# Neuroprotective effects of motherhood on brain function in late-life: a resting state fMRI study

**DOI:** 10.1101/2020.06.09.143511

**Authors:** Edwina R Orchard, Phillip GD Ward, Sidhant Chopra, Elsdon Storey, Gary F Egan, Sharna D Jamadar

## Abstract

The maternal brain undergoes structural and functional plasticity during pregnancy and the postpartum period. Little is known about functional plasticity outside caregiving-specific contexts, and whether changes persist across the lifespan. Structural neuroimaging studies suggest that parenthood may confer a protective effect against the ageing process, however it is unknown whether parenthood is associated with functional brain differences in late-life. We examined the relationship between resting state functional connectivity and number of children parented in 220 healthy older females (73.82±3.53years) and 252 healthy older males (73.95±3.50years). We compared patterns of resting state functional connectivity with three different models of age-related functional change to assess whether these effects may be functionally neuroprotective for the ageing human parental brain. No relationship between functional connectivity and number of children was obtained for males. For females, we found widespread decreasing functional connectivity with increasing number of children parented, with increased segregation between networks, decreased connectivity between hemispheres, and decreased connectivity between anterior and posterior regions. The patterns of functional connectivity related to the number of children an older woman has parented were in the opposite direction to those usually associated with age-related cognitive decline, suggesting that motherhood may be beneficial for brain function in late-life.

---

The transition to parenthood is associated with changes in brain structure and function across pregnancy and the post-partum period (Hoekzema et al., 2017; Hoekzema et al., 2020; Kim, 2016; Kim et al., 2010; Kim et al., 2014; Luders et al., 2018; Luders et al., 2020). While the physiological adaptations of pregnancy, birth and lactation are temporary (Heidemann & McClure, 2003), the environmental changes and challenges of parenthood last a lifetime: as children grow and develop, new challenges continue to arise. Increased environmental complexity is compounded with additional children, requiring simultaneous care across different stages of dependency. However, the lasting impact of parenthood on the human brain is poorly understood.

Recent evidence has shown that reproductive experience may result in lifelong changes to the structure of the human brain. In middle-aged women and men, ‘brain-age’ – the difference between chronological age and estimated age based on structural magnetic resonance imaging (MRI; Franke & Gaser, 2019) – is associated with the number of children the person has parented (de Lange et al., 2019). That is, parents showed “younger-looking” brains with increasing number of children, suggesting a neuroprotective effect of parenthood on *brain-age*. In addition, middle-aged parents have faster response times and fewer errors on visual memory tasks than childless men and women (Ning et al., 2020). In healthy older adults aged over 70-years, we reported that cortical thickness differs between parents and non-parents of both sexes (Orchard et al., 2020). In particular, for mothers, number of children was associated with increased cortical thickness in the parahippocampal gyrus and improved verbal memory performance, suggesting that motherhood confers a cognitive benefit into late-life. Taken together, current evidence is indicative of a life-long effect of parenthood on the human brain, which, at least in mothers, may be neuroprotective in ageing (de Lange et al., 2019; Ning et al., 2020; Orchard et al., 2020).

There have been many studies investigating functional brain changes in relation to parenthood. The majority of these studies have used functional MRI (fMRI) to examine parents’ brain response to infant pictures, movies, or sounds of infant cries (see Swain, Lorberbaum, Kose, & Strathearn, 2007 for review). These studies have predominantly examined first-time parents, usually mothers, in early parenthood (first few years post-partum). Mothers show increased activation of brain networks including the reward/motivation network, salience and fear networks, and theory of mind networks in response to images and videos of their own infant, compared to unfamiliar infant stimuli (Cardenas, Kujawa, & Humphreys, 2019; Leibenluft, Gobbini, Harrison, & Haxby, 2004; Lorberbaum et al., 2002; Naaz, Knight, & Depue, 2019). Little is known about the cumulative hormonal and environmental impact of having multiple children, or parental brain function outside the context of responses to infant stimuli.

Resting state functional connectivity indexes the intrinsic connectivity of the brain when it is not engaged in a specific task. As an index of the ‘baseline’ connectivity of the brain, any change in resting-state functional connectivity associated with parenthood suggests that parenting is associated with global, generalised changes, beyond responses to infant stimuli. There is emerging evidence to suggest that early parenthood is associated with such generalised changes in resting brain function. Dufford Erhart, and Kim (2019) found increasing functional connectivity between the amygdala, anterior cingulate, nucleus accumbens and cerebellum with increasing time since birth (parturition) in first-time mothers, from 0 to 10 months postpartum. Dufford et al. concluded that connectivity between reward/motivation and fear/salience networks may contribute to positive caregiving behaviours. Zheng et al. (2020) found reduced resting state functional connectivity within the default mode network in mothers at 3-months postpartum, compared to non-mothers. Zheng et al concluded that altered connectivity in the default mode network may reflect a neural adaptation to the demands of early motherhood, since mothers would be expected to have a stronger focus on infant-related responsibilities, and be reflective on the thoughts, feelings and needs of their children during the resting state.

In sum, there is emerging evidence to suggest that early parenthood is associated with changes in brain structure and function (e.g., Hoekzema et al., 2017; Zhang et al., 2020; Dufford et al., 2019), and that these changes may be neuroprotective against subsequent age-related brain changes in older adulthood (de Lange et al., 2019; Ning et al., 2020; Orchard et al., 2020). In the current study, we examined whether parenthood conveys resilience against age-related functional brain changes in humans using resting-state fMRI. Healthy ageing is associated with widespread changes in functional brain activity. Task-related brain activity of healthy older adults is associated with reduced hemispheric lateralisation (hemispheric asymmetry reductions in older adults; HAROLD effect; Cabeza, 2002) and a posterior-to-anterior shift in fMRI activity (PASA effect; Davis, Dennis, Daselaar, Fleck, & Cabeza, 2008) compared to younger adults. HAROLD and PASA effects reflect compensatory recruitment of additional regions to support behavioural and cognitive function (Cabeza et al., 2018; Festini et al., 2019). In resting-state fMRI, older adults show lower within-network connectivity (reduced network integration), and higher between-network connectivity (decreased network segregation) compared to younger adults (Chan, Park, Savalia, Petersen, & Wig, 2014; Jamadar, 2019; Wig, 2017).

To test the hypothesis that parenthood conveys a neuroprotective effect against age-related neural changes, we correlated whole-brain functional connectivity matrices of mothers and fathers with the number of children parented. We compared the obtained matrices to three models of age-related neural change: HAROLD, PASA, and network integration/segregation (Figure 1). Since the HAROLD and PASA models are most commonly studied using task-related fMRI, we operationalised these patterns based on connectivity matrices sorted by anatomy (i.e., sorted by hemisphere, then by lobe). We defined a HAROLD effect as increased functional connectivity between lateral hemispheres, evident in the connectivity matrix as stronger connections edges between nodes in the left and right hemispheres (Figure 1a). We defined the PASA effect as increased connectivity between anterior and posterior brain regions, evident as stronger connections between frontal and posterior brain regions in the connectivity matrix (Figure 1b). Lastly, the network integration/segregation pattern is defined as changes in connectivity patterns within, and between, known resting-state networks. Therefore to test this model, we sorted brain regions into canonical networks. The network integration/segregation pattern is evident as lower connectivity along the matrix diagonal (i.e., within-network connectivity) and higher connectivity off-diagonal (i.e., between-network connectivity Figure 1c). Importantly, because we hypothesise that parenthood confers resilience to the ageing process, we hypothesised that as the number of children parented increases, older adults would show *reduced* HAROLD, PASA and network integration/segregation effects. In other words, that as number of children parented increases, older adults will show decreased connectivity between hemispheres (HAROLD), between frontal and posterior regions (PASA), and between networks (integration/segregation). Such a pattern of results would be consistent with the argument that older parents have ‘younger-looking’ brains (de Lange et al., 2019; Ning et al., 2020).

**Figure 1:**
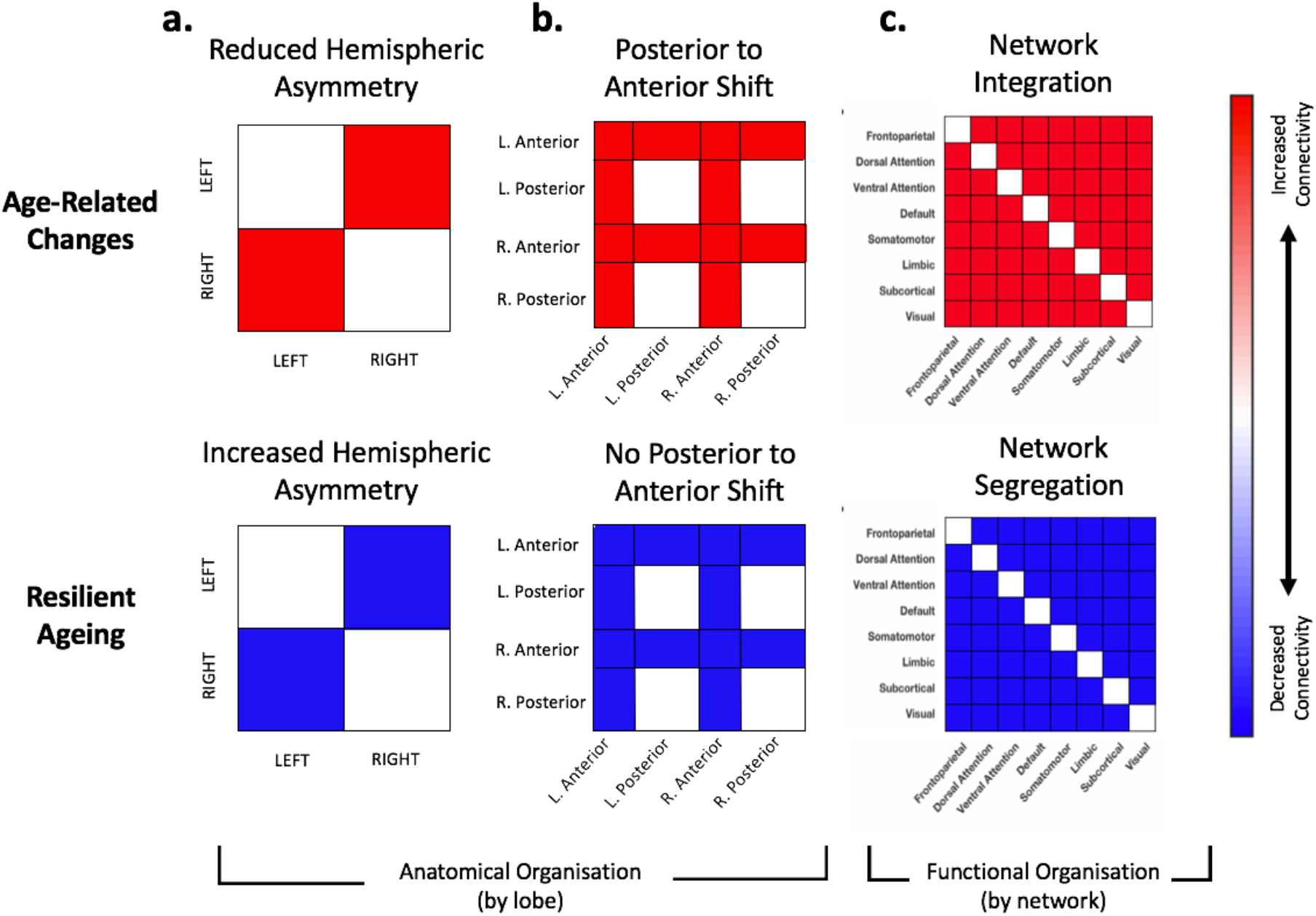
Cartoon schematics for three patterns of age-related functional brain changes in the late life parental brain. Red denotes increased resting state functional connectivity, and blue denotes decreased resting state functional connectivity. In order to assess hemispheric asymmetry and frontal compensation, regions are ordered according to anatomy (hemispheres and lobe assignment), whereas, assessing network integration/segregations requires a functional organisation (networks). **A. Hemispheric Asymmetry:** Cartoon schematic of the HAROLD effect (Hemispheric Asymmetry Reductions in Older Adults; (Cabeza, 2002)). Ageing is associated with reduced functional lateralisation. Therefore, results consistent with ageing in the HAROLD pattern would show *increased* functional connectivity between hemispheres (left-right/ right-left), shown in red. Decreased functional connectivity between hemispheres would be consistent with a reduced HAROLD effect (increased hemispheric asymmetry), indicative of resilient ageing, shown in blue. **B. Posterior to Anterior Shift:** Cartoon schematic of the PASA pattern (posterior-anterior shift in ageing; (Davis et al., 2008)). Ageing is associated with increased connectivity between anterior (prefrontal) regions and other areas (shown in red), consistent with a shift from posterior to anterior processing. Decreased functional connectivity between nodes of the frontal lobe and other regions would be consistent with a reduced PASA effect (less frontal compensation), indicative of resilient ageing (shown in blue). **C. Network Integration/Segregation:** Cartoon schematic of integration and segregation. Within-network connections lie on the diagonal (e.g. visual-visual), and between-network connections lie on the off-diagonal (e.g. visual-default mode). Ageing is associated with increased integration and decreased segregation of functional networks. Results consistent with age-related changes would show increased functional connectivity between networks (integration), shown in red. Decreased functional connectivity between networks (segregation) would therefore indicate resilient ageing, shown in blue.

## Methods

This data was acquired by the ASPREE Investigator Group, under the ASPREE-Neuro sub-study (S. A. Ward et al., 2017). Data are available on application from the ASPREE International Investigator Group (https://aspree.org). All methods for the ASPREE-Neuro clinical sub-trial were approved by the Monash University Human Research Ethics Committee, in accordance with the Australian National Statement on Ethical Conduct in Human Research (2007). This data was used to explore effects of parenthood on cortical thickness in late adulthood in our previous manuscript (Orchard et al., 2020).

### Participants

Baseline characteristics of the full ASPREE sample are reported in McNeil et al. (2017). We included data from baseline, prior to study medication (aspirin). Healthy adults in Australia were eligible to participate if aged 70-years and over, had no history of occlusive vascular disease, atrial fibrillation, cognitive impairment or disability; were not taking antithrombotic therapy, and did not have anaemia or high risk of major bleeding, or a diagnosis likely to cause death within five years. At study entry, each participant had a Modified Mini Mental Status Examination (3MS; Teng & Chui, 1987) score of at least 78/100, and were able to perform all six of the Katz activities of daily living (ADLS; Katz & Akpom, 1976).

MRI data from the ASPREE-Neuro sub-trial, *N*=472 participants aged 70-87 years (mean age [+/-stdev] = 73.9+/−3.5years) were used, comprising n=252 males (73.9 ±3.5 years), and n=220 females (73.8 ±3.5 years). As part of a health outcomes questionnaire, ASPREE-Neuro participants were asked “How many children do you have?”. More detailed parenthood data (e.g. biological/adopted children, grandparenthood, primary/secondary caregiver, absence from children, age at first/last pregnancy etc.) is not available. The number of children parented by males and females in our sample is shown in Table 1.

**Table 1:** Number of participants with 1-6 children for males and females.

|  | Mothers | Fathers |
| --- | --- | --- |
| 1 child | 19 | 16 |
| 2 children | 81 | 106 |
| 3 children | 71 | 88 |
| 4 children | 36 | 26 |
| 5 children | 8 | 11 |
| 6 children | 5 | 5 |
| **Total** | **220** | **252** |

### Image Acquisition and Preparation

All MRI scans were obtained on a 3T Siemens Skyra MR scanner (Erlangen, Germany) with a 20-channel head and neck coil at Monash Biomedical Imaging, Melbourne, Australia. Here, we used T1 and resting state functional magnetic resonance imaging (rsfMRI) data. T1-weighted magnetization-prepared rapid gradient-echo (MPRAGE) images were acquired (TR=2300ms, TE=2.13ms, TI=900ms, matrix size=256×240×192, bandwidth=230 Hz/pixel, 1×1×1mm^3^ voxels) and used for anatomical reference. Resting state data were obtained using multiband echo planar imaging (EPI) sequence (TR=754ms; multiband acceleration factor = 3; matrix size = 64×64; number of slices = 42; TE=21ms; number of time points=400; bandwidth=1474 Hz/pixel; 3-mm isotropic voxel (Xu et al., 2018)). Prior to functional data acquisition, a gradient echo-based field map was acquired to correct rsfMRI scan for geometric distortions. Additional sequences were acquired but not used in the present study; full details in Ward et al. (S. A. Ward et al., 2017).

T1- and T2-weighted MR images were segmented using the ‘recon-all’ function of FreeSurfer v5.3.0 (Dale, Fischl, & Sereno, 1999). Cortical thickness values were extracted for 41 regions for each hemisphere using the Desikan-Killiany Tourville atlas, producing 82 cortical thickness values in total (Desikan et al., 2006).

fMR images were corrected for geometry distortions (FUGUE) and the brain extracted (BET) (Smith, 2002). Intra-scan movements were corrected using 3dvolreg; both high frequencies (above 0.1Hz) and the temporal mean (and first- and second-order polynomials) were removed from each voxel’s time series using 3dTproject (AFNI; Cox, 1996). Filtered images were entered into a first-level independent component analysis with automatic estimation of the number of components using MELODIC (Beckmann, DeLuca, Devlin, & Smith, 2005). All the extracted ICA maps were then automatically labelled by FSL-FIX (Salimi-Khorshidi et al., 2014), which was previously manually trained on 25 random subjects. Temporal trends from noise-labelled ICA components were linearly regressed out of the 4D MR images using ordinary least squares (OLS) regression as implemented in FSL FIX. The cleaned images were then normalized to the MNI template (2mm isotropic resolution); the first volume of the EPI time series was registered to the T1-weighted image using linear registration (with 6 degrees of freedom). Each T1 was then non-linearly registered to the MNI template using the symmetric normalization algorithm in ANTs (Avants, Epstein, Grossman, & Gee, 2008). The brain was parcellated into 82 regions using the Desikan-Killiany-Tourville atlas (Desikan et al., 2006; Fischl et al., 2002) and then applied to the EPI. Finally, the normalized and cleaned file was smoothed with a 5mm FWHM Gaussian kernel.

### Data Analysis

#### Functional connectivity

fMRI timeseries were extracted from each of the 82 Desikan-Killiany-Tourville regions for each individual. An individual functional connectivity matrix was calculated as an 82×82 correlation matrix, formed by calculating temporal correlations (Pearson) between each pair of regions. A single entry in the correlation matrix is referred to as an ‘edge’ in the connectivity matrix; each region is referred to as a ‘node’.

Subsequent data analysis focused on testing the hypotheses of the effects of parenthood on functional connectivity. We examined the effects of parenthood for males and females separately, given the substantial differences in the experience of parenthood between the sexes. For example, women experience the hormonal effects of pregnancy, parturition, and breastfeeding in addition to the environmental effects of child-raising (which are experienced by both sexes, albeit to a different extent depending on caregiving roles within the family). Exploratory comparisons of connectivity between the sexes were not carried out, given recent evidence suggesting that BOLD signal intensity and functional connectivity, is confounded by haemoglobin differences between the sexes (Ward et al., 2020).

#### Effects of parenthood on functional connectivity

As the number of children parented is discrete, ordinal data, with a non-normal distribution, all analyses used were non-parametric. Mass univariate testing using a two-step procedure was performed to compute Spearman’s correlation between the number of children parented and functional connectivity at each edge. In the first step, we used a GLM to adjust functional connectivity values for the effects of age, education, haemoglobin (P. G. Ward et al., 2019), and cortical thickness (Damoiseaux, 2017) on parenthood-connectivity relationships. In the second step, we computed a Spearman’s correlation coefficient between the residualised functional connectivity and number of children.

For statistical inference and family-wise error correction, the Network Based Statistic (NBS) was used (Zalesky, Fornito, & Bullmore, 2010). The extent (number of edges) of any connected-components in the observed data that survived a predetermined primary component-forming threshold of *p*<0.01 were compared to a null-distribution of maximal component sizes from 10,000 permutations. At each permutation, the number of children was randomly shuffled between subjects, without replacement. Then, the same mass-univariate two-step procedure which was used to compute the correlation between functional connectivity and number of children parented in the observed data, was applied to each edge. This provided a null distribution of Spearman’s Rho (*ρ*) values at each edge, which were then used to obtain non-parametric edge-level *p*-values for the observed data. For component-level family-wise-error correction, each permutated 82 × 82 matrix of ρ values was thresholded at a critical ρ value equivalent to *p*<0.01. This critical value was determined by combining the ρ values from all permutations into one large null distribution and selecting the value at the 99^th^ percentile as the critical equivalent to *p*<0.01, which in the current study was ρ=.174. The extent of the largest connected component that survived this threshold at each permutation was recorded. This resulted in an FWE-corrected null distribution of maximum component sizes. Any connected component in the observed data that survived the primary threshold and was larger than the 99^th^ percentile (*i.e*. *p*< 0.01 FWE-corrected) was deemed statistically significant (Zalesky et al., 2010).

To examine whether the pattern of parenthood-connectivity was consistent with the three patterns of age-related functional brain changes, we created a matrix of significant edges, and ordered nodes in terms of their anatomical organisation (HAROLD and PASA) or functional networks (integration/segregation).

To quantify agreement with these models, we first defined the patterns of connectivity consistent with each model (Figure 1). We then calculated the number of observed edges consistent with each (i.e. inter-hemispheric connections, frontal lobe connections, between network connections). We then randomly permuted a random network of the same size 10,000 times and calculated the number of model-consistent edges in each permutation. Comparing the number of model-consistent edges between observed and permuted data allowed non-parametrically computation of significance, i.e., the likelihood of observing these patterns in networks with edges randomly distributed across the brain.

The HAROLD effect was defined as increased functional connectivity between lateral hemispheres, evident in the connectivity matrix as stronger connections edges between nodes in the left and right hemispheres (Figure 1a). To test the HAROLD model, the ROIs were sorted by hemisphere (left, right) then anatomical division (frontal, parietal, subcortical, temporal, occipital); and then calculated whether parenthood-related connectivity appears as increased functional connectivity between the left and right hemispheres (consistent with age-related change) or decreased functional connectivity between the left and right hemispheres (consistent with resilient ageing).

The PASA effect was defined as increased connectivity between anterior and posterior brain regions, evident as stronger connections between frontal and posterior brain regions in the connectivity matrix (Figure 1b). To test the PASA hypothesis, the ROIs were sorted by anatomical division (by hemisphere then lobe: frontal, parietal, subcortical, temporal, occipital) and calculated whether parenthood-related connectivity appears as increased connectivity between frontal and posterior brain regions (consistent with age-related change) or decreased functional connectivity between frontal and posterior brain regions (consistent with resilient ageing).

Lastly, the network integration/segregation pattern is defined by changes in connectivity patterns between regions (nodes) of canonical resting-state neural networks To test the network integration/segregation hypothesis, the ROIs were sorted by network (visual, somatomotor, dorsal attention, ventral attention, limbic, frontoparietal, default mode, and subcortical; as classified by Yeo et al. (2011), and calculated whether parenthood-functional connectivity relationships showed greater within- or between-network connectivity. Two networks are ‘segregated’ if they have low/negative between-network functional connectivity, as their patterns of activation are not highly correlated.

Additionally, we were interested in determining which networks showed the largest effect of parenthood. In order to do this, we first had to correct for network size bias. The ROIs are not evenly distributed between networks, such that some networks (e.g. default mode network) contain a higher number of nodes than other networks. As such, there is a higher probability that significant edges would occur within networks with more nodes by chance. This bias was accounted for by normalising the count of significant edges by the network capacity, i.e., the total number of possible edges of each network.

## Results

### Functional connectivity

We examined the relationship between late-life brain function and number of children parented in healthy older females and males (Figure 2). Visual inspection of Figure 2 suggests that males showed strong (i.e., average beta-values between −0.70 and 1.35) resting state functional connectivity within anatomical sub-divisions, particularly within the frontal, parietal and subcortical regions, both within and between hemispheres (Figure 2a). A number of long-range connections between anatomical sub-divisions were also evident, including fronto-parietal, parietal-subcortical and frontal-subcortical regional connectivity. These long-range connections were also evident in older females (Figure 2b), but were of smaller magnitude than in males. Visual inspection of the resting-state connectivity matrix for females, showed strong (i.e., average beta between −0.88 and 1.26) connectivity within subcortical regions, and strong connectivity between subcortical regions and frontal and parietal regions. Occipital regions showed negative (average beta between −0.67 and 0.00) functional connectivity with frontal, parietal, subcortical and temporal regions, both within and between hemispheres in both males and females.

**Figure 2:**
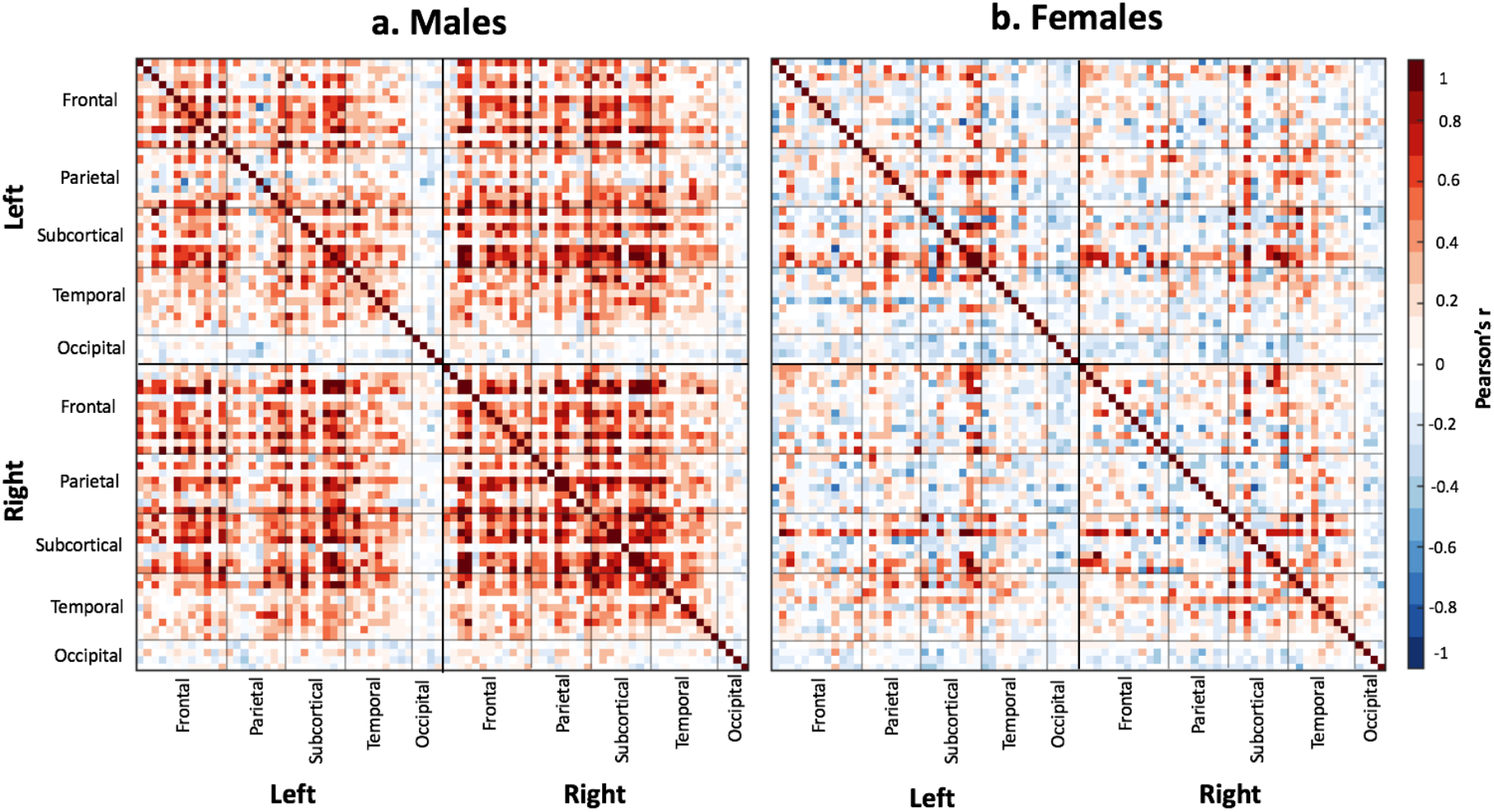
Mean functional connectivity for healthy older males (*N*=252) and females (*N*=220) who parented between one and six children. Each row and column of the 82×82 matrix represents a node (41 nodes in the left hemisphere and 41 in the right), organised by anatomical division (i.e., frontal, parietal, subcortical, temporal, occipital). Each cell represents a the functional connectivity between two nodes, after adjusting for age, education, haemoglobin and cortical thickness. Blue cells indicate negative edge weights and red cells indicate positive edge weights. The matrix has diagonal symmetry, such that the upper and lower triangles are mirrored. These matrices show the global functional connectivity for mothers and fathers, and do not reflect results of any significance tests.

### Effects of parenthood on functional connectivity

No significant connected component was found to be related to fatherhood for the men, so the following results pertain to the mothers only.

We obtained a 47-edge connected component related to the number of children a woman has mothered (1 to 6 children; *p*<.01 FWE-corrected). As a general observation, all edges in the network are negatively weighted, which indicates *decreasing* functional connectivity between nodes with increasing number of children.

### Hypothesis 1: Increased hemispheric asymmetry

Older mothers showed a pattern of increased hemispheric lateralisation with increasing number of children parented. Regions with high degree (number of edges; Table 2) consistent with a reduced HAROLD effect, include the bilateral frontal pole, bilateral inferior parietal lobule and bilateral posterior cingulate. Seventy percent of all observed edges (33/47) are between-hemisphere edges (left-right, or right-left connections); which is 1.4 times more edges between hemispheres than expected by chance (*p*<.001; Figure 3). This pattern of hemispheric lateralisation is in the opposite direction to that predicted by the HAROLD effect of age-related changes in fMRI activity.

**Table 2.**
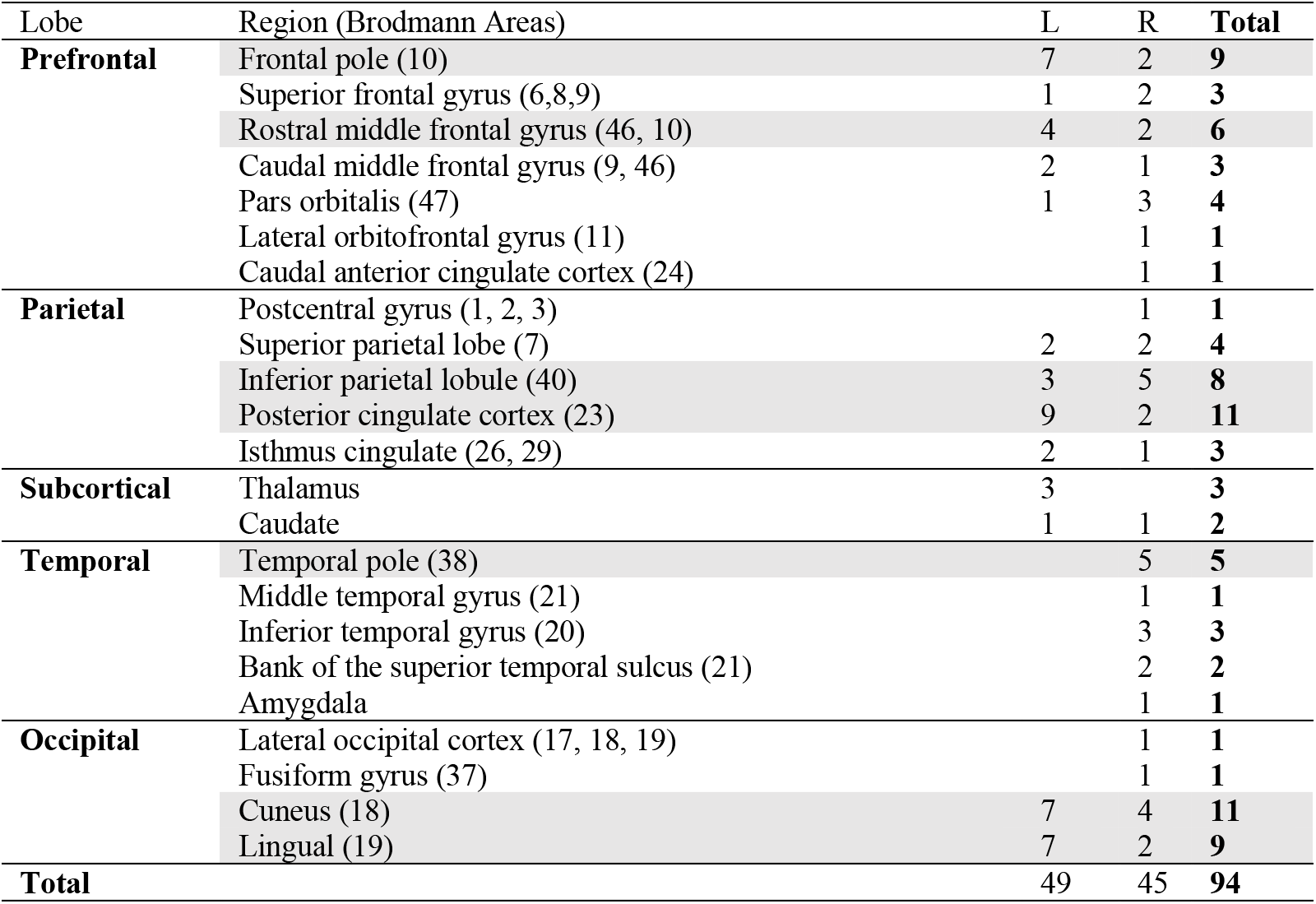
Degree (number of significant edges) for each node in the motherhood network. N.B. The motherhood network consists of 47 edges, but in calculating degree, each edge is counted twice (once for each node in the edge), so total degree is 94.

**Figure 3:**
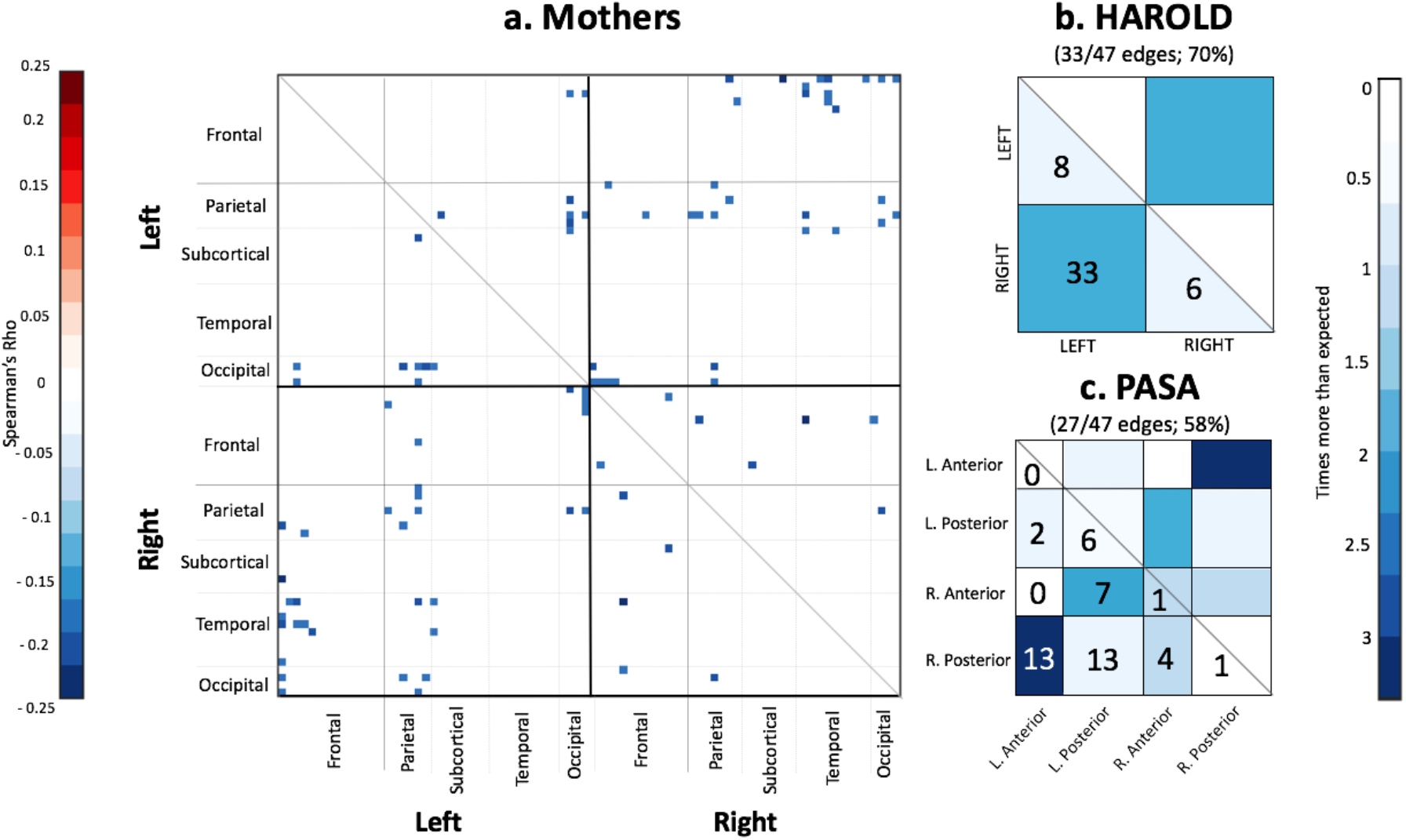
**A. Resting State Functional Connectivity for Mothers:** Adjacency matrix of pairwise correlations for the 47 edges associated with motherhood. The 82 ROIs are arranged by hemisphere then by anatomical division (frontal, parietal, subcortical, temporal, occipital) such that each row and column of the 82×82 matrix represents one node (41 nodes in the left hemisphere and 41 in the right). Each cell represents a significant edge between two nodes. The matrix has diagonal symmetry, such that the upper and lower triangles are mirrored, and each contains 47 edges. Blue cells signify negative edge weights and red cells signify positive edge weights. All edges related to motherhood are negatively weighted, showing decreasing functional connectivity between node pairs as number of children increases. **B. Schematic of results in relation to HAROLD (Hemispheric Asymmetry Reductions in Older Adults; (Cabeza, 2002)):** The blue quadrants highlight between-hemisphere connections (33 left-right and right-left edges), which is 1.4 times as many edges between hemispheres as would be expected by chance. The lighter coloured quadrants highlight within-hemisphere connections; left-left (8 edges; one third [0.34] as many edges as would be expected by chance) and right-right (6 edges; one quarter [0.25] of what would be expected by chance). Older mothers show decreased functional connectivity between hemispheres with increasing number of children. Seventy percent (33/47) of the observed edges are between-hemisphere connections. Decreasing functional connectivity between the left and right hemispheres indicates *increased* hemispheric asymmetry with increased number of children mothered. This pattern is consistent with a *reduced HAROLD effect*, indicative of resilient ageing in mothers. **C. Schematic of results in relation to PASA (Posterior to Anterior Shift in Ageing; (Davis et al., 2008)):** The blue bars highlight connections with the left and right prefrontal cortex, coloured to compare the observed edges with the number that would be expected by chance. For example, there are 3.2 times as many edges between left anterior (frontal) regions and right posterior regions than would be expected by chance. Sixty-four percent (30/47) of the observed edges show decreased functional connectivity between regions of the prefrontal cortex and other regions, indicating *less* frontal compensation with increasing number of children mothered. This pattern is consistent with a *reduced PASA effect*, indicative of resilient ageing.

### Hypothesis 2: Reduced posterior to anterior shift

Mothers showed a pattern of decreased frontal lobe integration with increasing number of children parented. Specifically, between the bilateral frontal pole, superior and middle frontal gyri and inferior frontal gyrus pars orbitalis, and posterior brain regions. Fifty eight percent of the observed edges (27/47) are between nodes in the prefrontal cortex and posterior brain areas; overall 1.7 times more than expected by chance (*p*=.001; Figure 3). This pattern is in the opposite direction to that predicted by the PASA model on age-related functional brain activity.

### Hypothesis 3: Increased network segregation

As the number of children increased, mothers had increased network segregation, with the majority (45 edges; 96%) of the negatively-weighted edges *between* networks (*p*=.002) compared to *within* networks (2 edges; 4%;). Network segregation was not evenly distributed across the brain. There were nine times more edges between the fronto-parietal network and dorsal attention network, seven times more edges between the fronto-parietal network and visual network, and four to five times more edges between the default mode network and visual network in the default mode network compared to what would be expected in a randomly distributed network of the same size (Figure 4).

**Figure 4:**
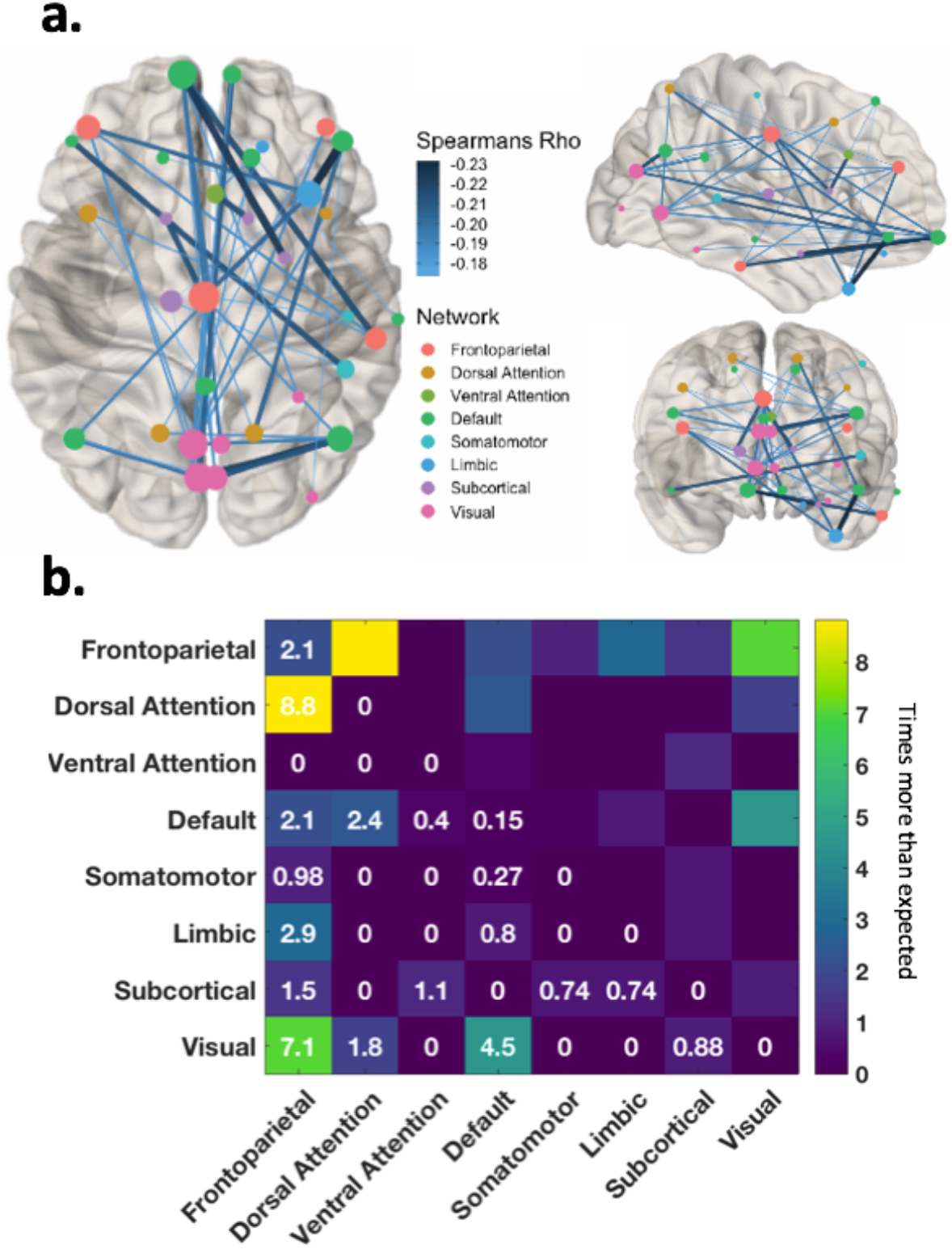
Functional connectivity within and between brain networks (i.e. frontoparietal, dorsal attention, ventral attention, default mode, somatomotor, limbic, subcortical and visual; as classified by Yeo et al. (Thomas Yeo et al., 2011)). **A**: MNI-space network graph of the edges associated with motherhood (left: axial; top: coronal; bottom: sagittal). The nodes are weighted by degree, such that nodes with a higher number of connections are larger. Edges are weighted by the strength of Spearman’s Rho, such that edges with stronger connectivity are darker and thicker **B**: **Segregation and Integration:** Functional connectivity within and between networks. Within-network connections lie on the diagonal (e.g. visual-visual), and between-network connections lie on the off-diagonal (e.g. visual-default mode). The numbers denote how many more edges are observed between networks, compared to what we would expect by chance, after adjusting for the capacity of the network, e.g we observed 7.1 times more edges between the frontoparietal network and the visual network than we would expect by chance. All edges related to motherhood are negatively weighted, therefore decreasing connectivity between networks (off-diagonal) indicates increased network segregation with increasing number of children mothered.

## Discussion

Our results show a relationship between number of children parented and functional connectivity in the late-life parental brain. In mothers, there was a significant component with 47 negatively-weighted edges, indicating decreased functional connectivity with increased number of children. No significant component was found between number of children and functional connectivity in fathers, therefore the discussion focuses on motherhood and the influence of parenting multiple children on brain function in the late-life maternal brain.

Previous studies have reported neuroprotective effects of parenthood on brain structure later in life in mothers (de Lange et al., 2019; Ning et al., 2020; Orchard et al., 2020), and fathers (Ning et al., 2020). Therefore, we compared resting-state connectivity in older parents to three patterns of age-related functional activity, to assess whether the number of children a woman has parented is related to patterns of resilient brain function in late-life. Here we show patterns of functional connectivity in older mothers that show: i) increased hemispheric lateralisation, ii) decreased frontal lobe integration, and iii) increased network segregation. These patterns are in the opposite direction to that predicted by three patterns of functional brain changes in ageing, the HAROLD and PASA models, and the network integration/segregation effect. That is, as the number of children parented increased, women’s resting-state brain activity became less similar to patterns of age-related functional brain change. Importantly, we do not suggest that mothers show *no* age-related brain changes, or that they are showing ‘reverse’ ageing effects, rather, that patterns of age-related changes in the brain are *less* evident in women with more children than those with fewer children. This is consistent with previous results suggesting mothers show “younger-looking” brains (e.g., de Lange et al., 2019). Taken together, these results suggest that motherhood confers functional neuroprotection for the ageing maternal brain.

The first pattern of age-related functional changes that we investigated was the Hemispheric Asymmetry Reductions in Older Adults (HAROLD; Cabeza, 2002) effect, which represents a pattern of reduced functional lateralisation in older adults compared to younger adults. We found a pattern of brain connectivity in the opposite direction to that predicted by the HAROLD model: mothers showed decreased connectivity between the left and right hemispheres with increasing number of children. In other words, mothers with more children have retained hemispheric asymmetry, consistent with a resilient pattern of brain ageing, suggesting a protective effect of reproductive experience on the function of the ageing maternal brain.

The second pattern of age-related functional changes that we studied was the posterior-to-anterior shift in ageing (PASA) effect (Davis et al., 2008). In task-related fMRI, the PASA effect has meta-analytic support (Li et al., 2015), and is interpreted as increased compensatory activation of the prefrontal cortex, however it is unclear if PASA occurs in response to varying task demands (Festini et al., 2019). Consistent with the overall pattern of results in this study, we found that the PASA effect reduced with increasing number of children a woman had parented, again suggesting that mothers display patterns of resilient brain function in late life.

For the third pattern of age-related functional changes, we examined patterns of network integration/segregation. In healthy ageing, functional brain networks become less segregated and more integrated, apparent as weaker connectivity within networks, and greater connectivity between networks (Chan et al., 2014). Network segregation is considered beneficial for brain function, as the networks are more functionally specialised, more easily adaptable to task demands, and more resilient (Wig, 2017). We found segregation between the frontoparietal network, the default mode network, the dorsal attention network, and the visual network. The frontoparietal network plays a central role in executive control and adaptive behaviour via flexible top-down modulation of other brain regions (Dixon et al., 2018). The frontoparietal network is flexibly co-activated with the default mode network and dorsal attention network, modulating introspection and self-referential processing (supported by the default mode network) or perceptual attention (supported by the dorsal attention network), depending on task demands (Dixon et al., 2018; Spreng, Sepulcre, Turner, Stevens, & Schacter, 2013). The task-negative default mode network and the task-positive dorsal attention network typically show anticorrelated activity in both resting-state and task-based functional connectivity, and this competitive relationship impacts behaviour and cognitive function. Here, we found that mothers with more children showed increased network segregation, with almost all of the negatively-weighted edges *between* networks, rather than *within* networks. This suggests that as the number of children an older woman has parented increases, the more similar the brain functions to that of a younger person’s brain. These results are consistent with a more flexible and resilient late-life maternal brain.

Results in the default mode network, specifically in the posterior cingulate, are also consistent with those previously found in the early postpartum period. Zheng et al. (2020) found reduced resting state functional connectivity within the default mode network in mothers at 3months postpartum, compared to non-mothers, using the posterior cingulate cortex as a seed. The posterior cingulate is involved in outward, preventative aspects of self-relevant thought, including duties and responsibilities to others (Johnson et al., 2006), as well as theory of mind abilities (Mars et al., 2012). The behaviours of early motherhood may be reflected by this functional adaptation, as motherhood requires a heightened focus on infant-related responsibilities, and more in tune with the thoughts, feelings and needs of their children during rest (Chase, Moses-Kolko, Zevallos, Wisner, & Phillips, 2014). Therefore, default mode network segregation may be a motherhood-related functional brain change that is altered in the early postpartum period (3-4months postpartum) and endures across the life span.

The current functional connectivity results are in line with previous reports of structural brain changes in the ageing parental brain. In our previous study using the same sample, we found that the number of children an older woman mothered was associated with thicker cortical grey matter in the parahippocampal gyrus (Orchard et al., 2020). The parahippocampal gyrus is involved in memory consolidation and cohesion (Köhler et al., 1998), and atrophy in the parahippocampal gyrus plays a key role in age-related memory decline (Burgmans et al., 2011). Furthermore, we found that verbal memory performance improved with increasing number of children mothered. A positive association between number of children parented and cortical thickness in a region associated with memory suggests that motherhood confers a cognitive benefit into late-life. This is consistent with the finding that middle-aged women show “younger” brain-age with increasing number of children (de Lange et al., 2019; Ning et al., 2020). Taken together, the results of the current and previous neuroimaging studies of the effects of parenthood on the ageing brain are consistent with a neuroprotective effect of motherhood. That is, that the effects of motherhood on the structure and function of the brain are both beneficial and long-lasting, persisting beyond the first few postpartum years.

While the current pattern of results are compatible with previous studies of the effects of parenthood on the structure of the late-life brain, the pattern of results obtained in the immediate post-partum period are less clear (reviewed in Orchard et al., 2020). Some studies show that early motherhood is associated with reduced grey matter in pregnancy (Hoekzema et al., 2017), whereas others show that the postpartum period is associated with increases in grey matter (Kim, Dufford, & Tribble, 2018; Kim et al., 2010; Luders et al., 2020). Regardless of the direction of the effect, interpreting changes in cortical thickness and its relationship to improved or impaired functional outcomes is not straightforward. In the cognitive ageing field, reduced cortical thickness is usually interpreted as a negative consequence of ageing; with thinner cortex associated with poorer functional outcomes (e.g. Fjell et al., 2006; Storsve et al., 2014). In contrast, in the parenthood field, both reduced (e.g. Hoekzema et al., 2017), and increased grey matter (e.g. Kim et al., 2018; Kim et al., 2010; Luders et al., 2020) has been interpreted as a positive outcome. By comparison, patterns of resting-state functional connectivity associated with parenthood are more easily interpretable. Our results indicate that the functional brain activity of older mothers with more children resembles “younger looking” patterns of brain activity.

We argue that late-life functional brain differences associated with parenthood are likely triggered by endocrine changes of pregnancy, birth and lactation, and are maintained across the lifespan by the ongoing environmental complexity of parenthood (Orchard et al., 2020). While the dramatic hormonal shifts of pregnancy are likely to play an initial role in maternal brain plasticity, hormone levels return to a pre-pregnancy state after birth (Tulchinsky, Hobel, Yeager, & Marshall, 1972), and most biological changes are resolved in the first year, i.e., uterus size, lactation, etc. (Henry et al., 2016). In our sample, the hormonal exposure of pregnancy likely occurred three or more decades prior to testing. In contrast, the environmental complexity of parenthood endures throughout the entire parenting experience, possibly throughout the life-span of a parent, compounding with increasing number of children. This environmental complexity is consistent with the conceptualisation of caregiving as a dynamic, learning experience in humans (Anderson, 2011; Parsons et al., 2017), and as a period of plasticity (Kim, 2016; Kim, Strathearn, & Swain, 2016). We speculate, therefore, that motherhood may contribute to a person’s psychosocial or cognitive reserve (Cabeza et al., 2018; Stern et al., 2018), by increasing lifetime cognitive and social demands. Since resting-state designs cannot directly test whether differential brain activity is associated with improved cognitive function, we cannot decisively conclude that motherhood contributes to cognitive reserve. Future studies should examine whether motherhood contributes to cognitive reserve by using task-based cognitive neuroimaging designs that manipulate task difficulty (cf. Cabeza et al., 2018; Festini et al., 2019).

Our results suggest that high parity (number of children) is neuroprotective for patterns of brain function in mothers. However the relationship between motherhood and late-life health is more complex than ‘having more children is good for you’. Higher parity is associated with lower levels of education (a proxy for socioeconomic status; Gray & Evans, 2017; Huber et al., 2010). High parity is also associated with poorer late-life health, including increased late-life disability (Akin et al., 2010; Hank et al., 2010), obesity (Heliövaara & Aromaa, 1981), and increased cardiovascular risk in late life (Rich-Edwards et al., 2014). However, the relationship between parity and late life health is somewhat paradoxical: while parity is associated with higher risk of cardiovascular disease, stroke, and late-life vascular comorbidities, parity may also have a significant protective effect against cardiovascular disease *mortality* (Ritzel et al., 2017). In other words, while parity is associated with increased risk of developing those conditions, there appears to be a concomitant decreased risk of dying from those conditions. This paradox is also seen in the relationship between motherhood and Alzheimer’s disease, where women with more children have higher rates of Alzheimer’s disease pathology (Colucci et al., 2006), and show earlier disease onset (Sobow et al., 2004), but other evidence shows that more months of pregnancy over the lifetime is associated with reduced risk of Alzheimer’s disease (Fox et al., 2013). In sum, it seems that while motherhood and parity may be associated with negative physical health factors, these do not necessarily lead to the expected changes in brain health. This may be because of differential exposure to oestrogen, immunosuppression during pregnancy, cortisol, and variation in maternal hypothalamic-pituitary-adrenal (HPA) axis activity across the lifespan (Ritzel et al., 2017).

Here, the functional brain changes associated with parenthood were only evident in mothers. This result may reflect obvious biological differences between mothers and fathers (e.g. pregnancy, birth, breast-feeding), or differences in care-giving responsibilities between the sexes. Given the age demographics of our sample, it is likely that our sample were predominantly in ‘traditional’ care-giving arrangements, with primary care-giving mothers and ‘bread-winning’ fathers (Broomhill & Sharp, 2005) and that this may have limited the cumulative impact of parenthood for the males in this cohort. Although we did not obtain a relationship between parenthood and functional connectivity in fathers, this null result should be interpreted with care, particularly as there are other factors (e.g. pregnancy-induced effects, attachment, time spent in care) that are relevant for interpreting differing effects of parenthood on mothers and fathers, and as effects of fatherhood on late-life brain structure have been found in other large studies (Ning et al., 2020; Orchard et al., 2020).

The results of this study should be interpreted in context of the analytic framework. In the current study, the relationship between number of children parented and functional connectivity was obtained using Spearman’s rank-order correlation, which does not assume linearity, and instead measures the strength and direction of a monotonic relationship. However, the relationship may not be linear. De Lange et al. (2019) reported a modest non-linear relationship between brain structure and number of children parented in middle-aged women; however, this non-linearity was not replicated in their follow-up study, which found a linear relationship in the same sample (De Lange et al., 2020). Future studies should explore the potential for non-linear associations between parenthood and brain structure in middle-and late-life. Furthermore, the analysis reported here excluded participants who were childless (nulliparous), since nulliparous people cannot serve as an adequate ‘baseline’ or ‘control’ to which parental brains can be compared. Since having children was the Australian societal norm for people in this age group (approx. 90% of Australian women had children; Rowland, 1998), it raises the question of whether these women were child-free involuntarily. Infertility may be an indicator of poorer health more generally, and is associated with higher mortality in middle age (Grundy & Kravdal, 2008; Harville et al., 2018). In light of these considerations, we excluded nulliparous participants.

The primary limitation of this study is that the parenthood experiences of our cohort are minimally phenotyped. The data used in this study were acquired as part of a larger study with broader aims. Detailed demographic information related to parenthood was not obtained from the participants, and so information about the parenting experiences is not available. Many of these unknown factors, including birth and feeding methods, hormone levels, and parental sensitivity, potentially impact maternal neuroplasticity (Kim et al., 2016). We were also unable to control for pregnancies that did not result in live birth and/or parenthood experience (miscarriage, termination, still-birth, infant loss, or adoption), or exposure to child-rearing not associated with parenthood (e.g., grandparenthood, baby-sitting). Therefore, we are not able to confirm that some of our participants did not experience additional exposure to pregnancy hormones or interactions with children not associated with parenthood. It is also important to note that the Australian experience of parenthood has changed substantially since our participants became parents: for example, in the late 1960’s/early 1970’s, adoption rates were much higher (9,798 in 1971-72 vs. 310 in 2018-19; Australian Bureau of Statistics, 1998; Australian Institute of Health & Welfare, 2019), and forced adoption common practice for unmarried women and Australian Aboriginal and Torres Strait Islander people (Higgins, 2012). With the sparse information on parenthood, we are unable to control for these factors. Notably, *all* existing studies of the influence of parenthood on the late life brain are limited by the sparse information on parenting in their sample (de Lange et al., 2019; Ning et al., 2020; Orchard et al., 2020). This limitation reflects the fact that existing studies have reported the effects of parenthood on brain structure and function using large datasets acquired for other purposes (here, to examine the effects of aspirin exposure on brain structure; de Lange et al. (2019) and Ning et al. (2020) used data from the UK BioBank). Furthermore, it is important to note that reproductive experience is related to other traits which impact brain health in human ageing, including exposure to sex steroid hormones. Some of these traits are heritable (e.g. age at menarche and menopause). In the absence of these data, we cannot rule out the possibility that the observed relationships between functional connectivity and number of children parented are explained by some of these uncollected variables. Future studies should control for these factors, as well as examine the potential effects of grandparenthood and other opportunities for interaction with children during later life, with more richly-phenotyped parenting data. Lastly, the use of a cross-sectional design can only provide evidence of parenthood-related *differences* between groups, not *changes* associated with parenthood, which can only be established using longitudinal designs (Rugg et al. 2016).

Taken together, the resting state functional connectivity patterns shown in older mothers were in the opposite direction to three separate patterns of age-related functional changes, consistent with a neuroprotective effect of motherhood on the ageing human maternal brain. This finding shows patterns of resilient brain function in the ageing maternal brain across hemispheres, lobes, functional networks, and specific regions. This interpretation is in line with previous literature in humans and rodents, and paints a consistent picture of motherhood-related resilience in human ageing, and suggests functional effects in the human brain that are both life-long and beneficial.

## Acknowledgements

ERO, PGDW, SDJ, & GFE are supported by the ARC Centre of Excellence for Integrative Brain Function (CE140100007). SDJ is supported by an ARC Discovery Early Career Researcher Award (DE150100406) and National Health and Medical Research Council Fellowship (APP1174164). The authors acknowledge the assistance of Thomas Close, Linden Parkes and Francesco Sforrazini for their analytical support. The Corresponding Author is Sharna D Jamadar, 770 Blackburn Rd, Melbourne Vic, 3800, Australia.

The data in this manuscript were used under license from the ASPREE and ASPREE-Neuro studies (https://aspree.org). ASPREE-Neuro was supported by an Australian National Health and Medical Research Council Project Grant (APP1086188). ASPREE was supported by the National Institutes of Health (grant number U01AG029824); the National Health and Medical Research Council of Australia (grant numbers 334047, 1127060); Monash University (Australia) and the Victorian Cancer Agency (Australia). The Principal ASPREE study is registered with the International Standardized Randomized Controlled Trials Register, ASPirin in Reducing Events in the Elderly, Number: ISRCTN83772183 and clinicaltrials.gov number NCT01038583. ASPREE-Neuro trial is registered with Australian New Zealand Clinical Trials Registry ACTRN12613001313729.

## Competing Interests

The authors declare that there are no conflicts of interest.

## Author Contributions

ERO, SDJ & PGDW designed the research question; ERO, SC, PGDW conducted the analysis, ERO wrote the first draft of the paper, with revisions and supervision provided by SDJ, PGDW & GFE. ERO & SDJ obtained approval for use of ASPREE-Neuro data. ES & GFE sourced funding for this work. All authors have contributed to and approved this work.

## Notes

### Competing Interest Statement

The authors have declared no competing interest.

